# A targeted LC MS/MS assay of a health surveillance panel and its application to chronic kidney disease

**DOI:** 10.1101/2025.03.14.643399

**Authors:** Qin Fu, Casey Johnson, Lesley A. Inker, Jennifer E. Van Eyk, the CRIC Study Investigators

**Author notes:** Corresponding author, Jennifer Van Eyk, Advanced Clinical Biosystems Research Institute, Smidt Heart Institute, Cedars-Sinai Medical Center, 127 S. San Vicente Blvd., Advanced Health Science Pavilion A9307, Los Angeles, CA 90048.

## Abstract

Robust and reproducible assays capable of specific and quantitative monitoring of multiple biologically important proteins amongst the thousands of human plasma proteins can potentially be used to distinguish health versus disease. In this study, we established an LC-MS assay to monitor a Health Surveillance Panel (HSP) comprising 60 circulating plasma proteins selected based on their broad biological functions and assay performance. Plasma samples were prepared for proteomic analysis in an automated process. A scheduled LC-MRM assay with a 30-minute 5% - 35% acetonitrile gradient and 50.5 minutes of total run time was used to quantify the 60 endogenous proteins by monitoring 364 transitions from 117 proteotypic peptides along with their stable isotopic labeled standard peptides in a single assay. For each proteotypic peptide, we selected a quantifier ion and at least two qualifier ions. The quantifier ions have a linear response over a 100-fold range, and the peak area ratios of the three peptide ions were consistent. As proof of concept, we evaluated the performance of our HSP assay in a case-control study of progressive chronic kidney disease (CKD). Reduced plasma concentrations of alpha-2-antiplasmin, antithrombin-III, and immunoglobulin heavy constant alpha 1 correlated with CKD indicated by reduced GFR with p values < 0.05. These results demonstrate that the HSP proteins can be accurately and reproducibly quantified with a high-quality multiplexed MRM assay and the HSP assay can detect disease-associated differences.

## INTRODUCTION

Substantial progress has been made in quantifying proteins using targeted LC-MS quantification of proteotypic peptides unique to a protein of interest (1). Multiple reaction monitoring (MRM), also known as selected reaction monitoring (SRM), is a targeted and multiplexed MS workflow where at least one signature (proteotypic) peptide is quantified for each protein along with multiple fragment ions for each peptide (2, 3). Reliable MRM assays are an essential step in the translation of putative biomarkers from discovery into clinically applicable MS-based assays (4, 5). An MRM assay is considered reliable if the quantification of each target protein has assured repeatability, reproducibility, precision, and accuracy. This ensures that reliable results are obtained, which is needed when comparing biomarker expression between diseased and control groups (6, 7).

Plasma is a readily accessible biofluid commonly used for clinical research and clinical tests (8–10). It contains thousands of proteins and contacts virtually all cells in the human body. Plasma contributes to immunity, blood clotting, maintaining blood pressure, hemostasis, pH balance, and many other functions (11). Given its central role in human physiology, quantifying plasma protein concentrations should provide clinically useful information that reflects an individual’s health status (12–14).

Most FDA-approved plasma protein tests use single analyte sandwich immunoassays for quantification, rendering it a daunting task to monitor scores of proteins. However, MS-based assays can simultaneously detect multiple analytes in less time than a traditional ELISA. Adding stable isotopic labeled (SIL) peptide standards to an MS assay provides for unambiguous peptide identification and serves as an internal reference to increase quantitative accuracy while monitoring the performance of the MS system.

We propose that a subset of plasma proteins can be selected based on their known function and/or disease association to represent the overall health of an individual and assess their pathological state. This paper describes the development and performance of a scheduled MRM assay targeting a health surveillance panel (HSP) comprising 60 disease-associated plasma proteins. We propose that the HSP panel will be a valuable tool to establish associations between health status and the progression of disease.

As proof of concept, we tested the ability of the HSP assay to predict the progression of chronic kidney disease (CKD). Specifically, plasma was obtained from individuals who developed CKD at baseline when their glomerular filtration rate (GFR) was in the normal range and 3 years later when their disease had rapidly progressed and these samples were compared to age and sex matched healthy individuals with stable GFR over the same time period. Comparison of protein differences at the two time points revealed that reduced plasma concentrations of alpha-2-antiplasmin, antithrombin-III, and immunoglobulin heavy constant alpha 1 was correlated with CKD progression.

## EXPERIMENTAL PROCEDURES

### Chemicals, supplies and human plasma

Tris-(2-carboxyethyl)-phosphine (TCEP), methyl methanethiosulfate (MMTS), formic acid (FA),] 99.5+%, HPLC water, and acetonitrile (ACN) were purchased from Thermo Fisher Scientific (Waltham, MA USA). Octyl-beta-glucopyranoside (OGS), Tris-(2-carboxyethyl)-phosphine (TCEP), Tris base and CaCl_2_ were purchased from Sigma-Aldrich, Inc. Xbridge Peptide BEH30 C18 2.1mm x 100mm 3.5µm columns were purchased from Waters (Milford, MA USA). Trypsin treated with L-1-p-Tosylamido-2-phenylethyl chloromethyl ketone (TPCK) in 500 µg vials was purchased from SCIEX (Redwood City, CA USA). Tips for the Biomek i7 workstation and 1 mL deep well plates were purchased from Beckman Coulter (Indianapolis, IN USA). Pooled EDTA human plasma from healthy individuals was purchased from BioIVT (Westbury, NY USA).

### Stable isotopically labeled internal standard peptide mixture

Stable isotope labeled (SIL) peptides were synthesized and purified to >90% by HPLC by Synpel Chemical (Prague, Czech Republic) from SIL arginine (^13^C_6_,^15^N_4_) and SIL lysine (^13^C_6_,^15^N_2_) obtained from Cambridge Isotopes Laboratories Inc (Tewksbury, MA USA).

Automated peptide tuning was performed on a QTRAP 6500 with SIL peptides diluted in 20% ACN, 0.1% FA in HPLC water. Discovery Quant Software (Sciex) was used for automated tuning to obtain optimized multiple reaction monitoring (MRM) transitions and corresponding MS parameters: DP (volts), EP (volts), CE (volts) and CXP (volts). Well-behaved SIL peptides were mixed to create a SIL pool. The amount of each SIL peptide in the pool was titrated to yield a light endogenous peptide to heavy isotope peptide ratio of 0.1 – 10. The SIL peptides were aliquoted, dried in a speed vac, and stored at -80°C. For MRM assays, SIL peptides were resuspend in 0.1% FA. Each 10 μl sample injection contained 3 μg digested plasma and a 1X SIL peptide mixture in 2% ACN in 0.1%FA.

### Automated plasma sample preparation

Human plasma samples were prepared on a Beckman Coulter (Brea, CA USA) i7 Hybrid Workstation with a Dual-Multichannel head (a 96-Channel head and a Span-8 Pod). The digestion protocol was described previously (15, 16). Samples were loaded into a deep 96-well plate (Beckman Coulter) with a single channel pipette. All other liquid transfers were performed on the i7 Workstation operated with Biomek software version 5 (Beckman Coulter).

Typsin digestion was performed in deep 96-well plates, which were shaken at 1000 RPM for 15 seconds after addition of each reagent. Plasma 5 µl and 42.5 µl of mix 1 (2% OGS, 6 mM TCEP, 3mM CaCl2 and 75 mM Tris pH 8.2) were incubated with shaking (1000 RPM) at 60°C for one hour in the INHECO incubator. Next, 2.5 µL MMTS (200 mM) was added, and the plate was shaken for 10 minutes at 1000 RPM. Finally, 10 µL trypsin (Sciex) in 0.1% FA was added, and the plate was incubated and shaken at 1000 RPM for 2 hours at 43° C followed by addition of 10 µL of 10% FA to quench the reaction. The plate was centrifuged at 3400 RPM for 5 minutes at 4°C. 4 µL (∼30 µg) of digested plasma was transferred to a 96 well autosampler plate. Then, 90 µL of pooled SIL mixture (10 SIL unit in 2.2% of ACN and 0.1% FA) was added to each well. The plate was gently mixed at room temperature for one minute at 1000 RPM and then to loaded into the LC autosampler.

### Scheduled MRM assay

Tryptic peptides were mixed with SIL standard peptides in mobile phase A (2% ACN, 98% water 0.1% FA) and then analyzed on a Prominence UFLCXR HPLC system consisting of two LC-20ADXR pumps, a CTO-20AC controller, a CBM-20A light column oven, and a SIL-20 ACXR autosampler (Shimadzu, Japan) coupled to a QTRAP® 6500 triple quadrupole mass spectrometer (SCIEX, Framingham, MA) with a Turbo V Ion source. Analyst® software (version 1.6.3) was used to control the LC-MS/MS system and for data acquisition.

Mobile phase A consisted of 2% ACN, 98% HPLC-grade water, and 0.1% FA; and mobile phase B of 95% ACN, 5% HPLC-grade water, and 0.1% FA. The flow rate was 250 µL/min. For each run, 10 μl was injected onto a C18 XBridge Peptide BEH C18 analytical column (100 mm, 2.1 mm ID, 3.5 μm particle size, Waters, Milford, MA). The column temperature was set at 40 °C. After loading the diluted digest (equivalent to 0.05 µL plasma, or 3 µg protein and 1 unit of pooled SIL peptides), the column was equilibrated with 5% mobile phase B for 5 minutes.

Peptides were then eluted over 30 minutes with a linear 5% to 35% gradient of mobile phase B. The column was washed with 98% mobile phase B for 10 minutes and then returned to 5% mobile phase B for 5 minutes before loading the next sample. A two-phase switching valve was used to divert the post-column eluent to waste before it entered the ion source. MRM data were processed using MultiQuant™ 3.1 Software (SCIEX) for peak selection and integration and analyzed by comparing the intensity of sample peaks to SIL peaks.

Methods for the initial detection of potential HSP proteins by DIA (17) and DDA (18) have been previously described.

### Experimental design and statistical rationale

The CDK cohort comprised 32 participants from the Chronic Renal Insufficiency Cohort Study (CRIC) of the CKD BioCon Phase 1 of Biomarkers Consortium (19). The study design is shown in Figure S1. Cases were defined as those with a rapid decline in GFR (GFR slope >5 ml/min/year, total decline of at least 30 ml/min/1.73 m^2^) with a follow-up of at least 3 years. Controls were defined as those with a stable GFR (a decline of < 1 ml/min/year over at least 3 years). Cases and controls were matched based on age, sex, race, and diabetes status (Table 1). Plasma samples from baseline and at three years were aliquoted and stored at -80°C prior to sample preparation.

**Table 1.**
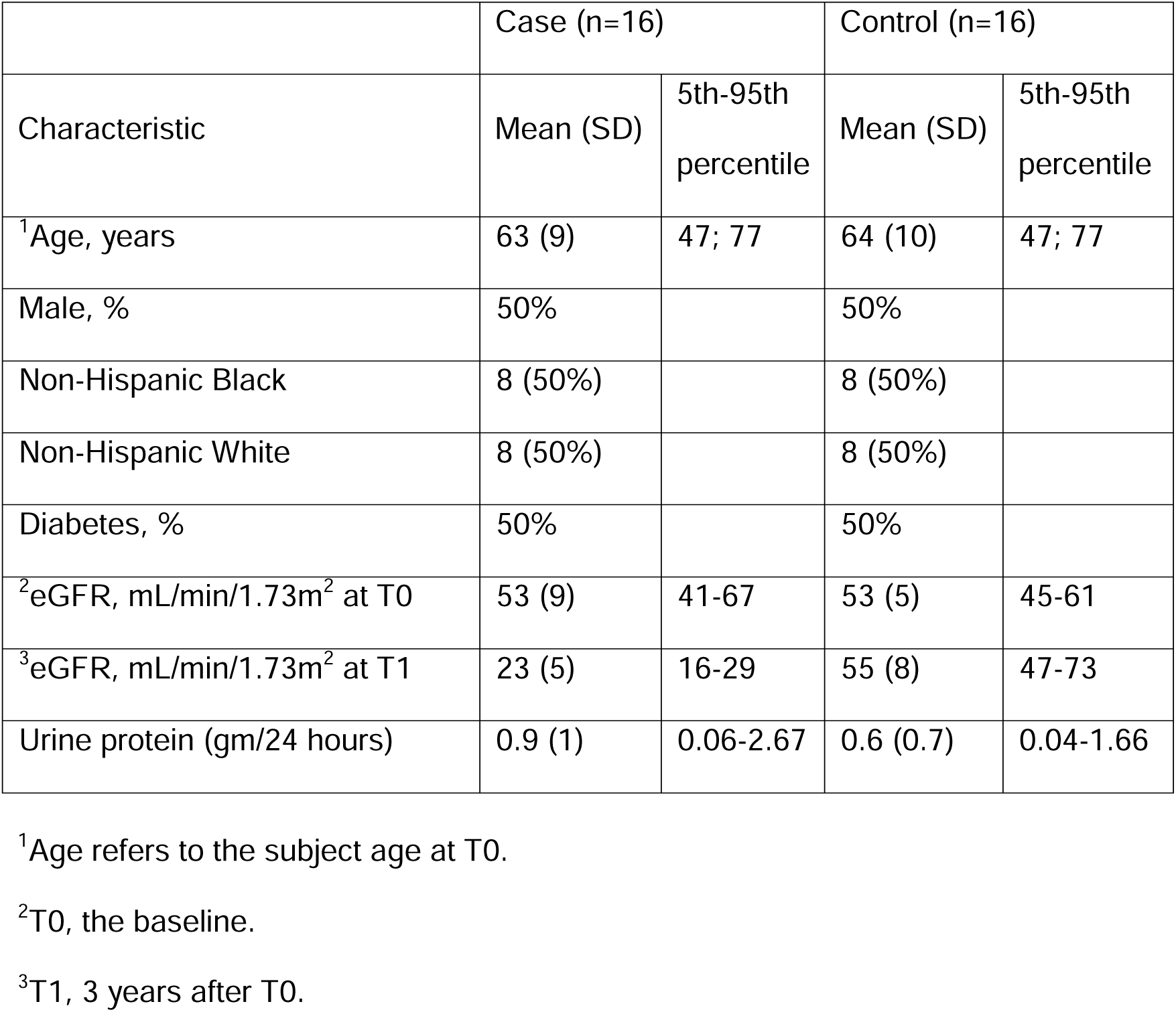
Baseline characteristics of individuals in the CRIC case-control cohort.

Differences between cases and controls at both time points were evaluated using a student’s t-test. Differences within cases or controls across time points were analyzed using a paired-t test. P-values <0.05 indicated a statistically significant difference.

All subjects gave their informed consent for inclusion before they participated in the Chronic Renal Insufficiency Cohort Study. The study was conducted in accordance with the Declaration of Helsinki, and the protocol was approved by the Institutional Review Board at each participating CRIC site.

## RESULTS

### Health Surveillance Panel

Plasma proteins for the HSP were selected based on a combination of literature search and in-house MS analysis of human plasma. Among the 60 analytes, 25 are currently measured in clinical laboratories with assays that have been cleared or approved by the FDA (Table S1).

The 60 HSP analytes were queried through the GeneALaCart batch query engine of GeneCards (20) and found to be associated with a wide variety of diseases including diabetes, heart diseases, kidney diseases, and immune system-related diseases and various biological processes including the innate immune response, platelet degranulation, complement activation, and endopeptidase activity (Figure 1, Table S2). The functions, biological processes, and localizations of the HSP proteins are similarly divers (Table S3).

**Figure 1.**
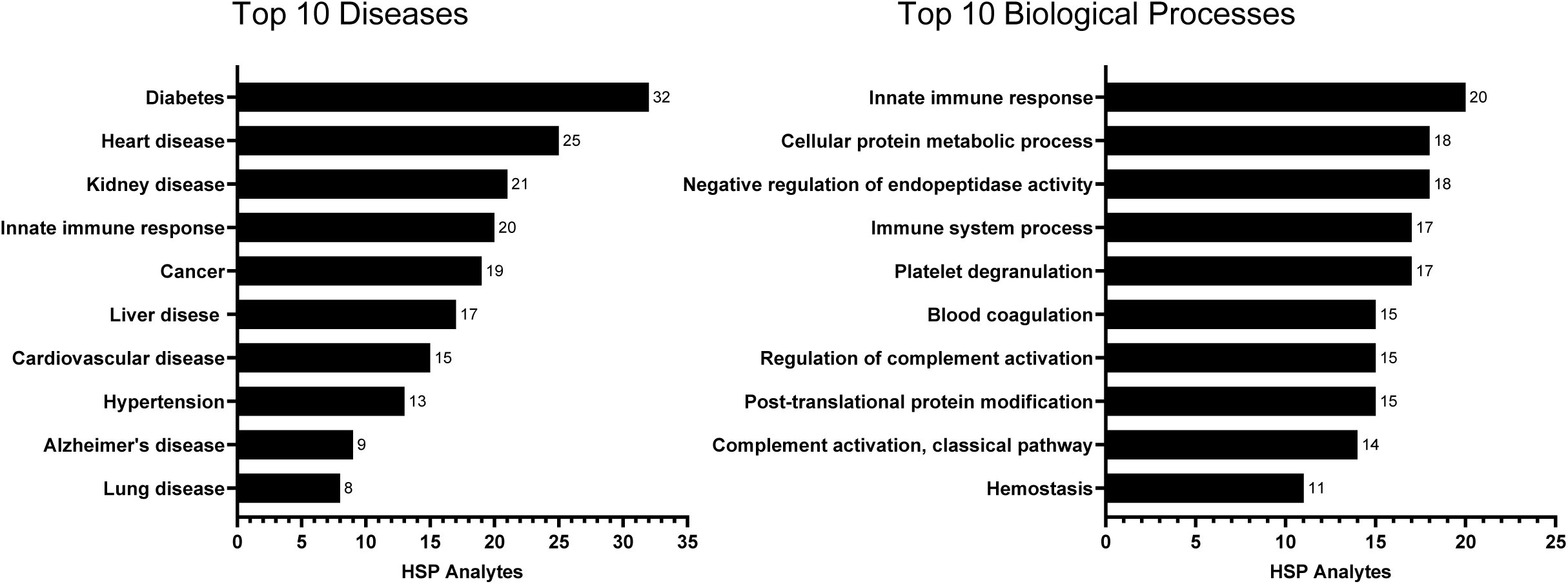
Examples of diseases and biological processes associated with HSP proteins, as indicated by GeneCards.^#^

### Peptide selection and MRM assay development

Quantifiable proteotypic peptides were selected for each protein using the workflow outlined in Figure 2. An *in silico* trypsin digest of the 60 HSP proteins yielded 2173 theoretical peptides having lengths of 6-30 amino acids. We evaluated 120 stable isotopic labeled (SIL) peptides with sequences that were proteotypic, readily detectable by MS of normal human plasma, free of cysteine or methionine residues, and representative of the 60 HSP proteins. The final scheduled MRM assay targeted 117 pairs of endogenous and SIL peptides by monitoring 364 pairs of native and heavy isotope standard transitions (Table S4).

**Figure 2.**
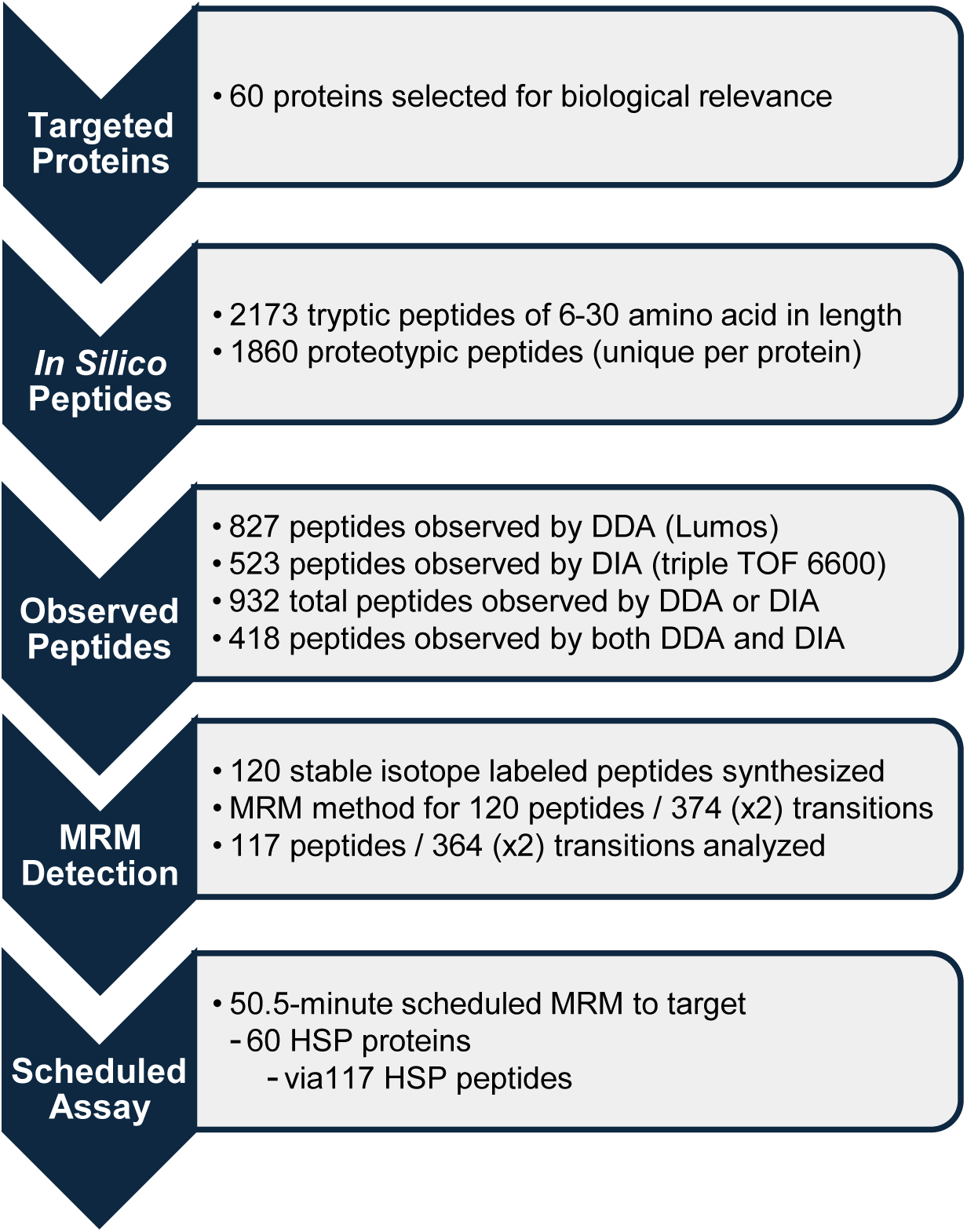
Process for developing the HSP assay.

### Peptide ion ratios

Ratios of the peak areas of two transition ions from the same peptide precursor (e.g. a high-intensity quantifier ion and a lower-intensity qualifier ion) are often monitored as a quality control step to evaluate the specificity of targeted LC-MSMS analyses (21, 22). In our multiplex MRM method, each peptide was measured by tracking at least three transitions: minimally a quantifier and two qualifiers (Table S5). We determined peptide ion ratios for SIL peptides spiked into 18 control plasma samples (Figure 3). For the 117 SIL peptides, the mean ion ratio of the first qualifier peptide (qualifier 1)/quantifier was 0.552 and the mean percent coefficient of variance (%CV) was 8.0%. The mean ion ratio of the second qualifier peptide (qualifier 2)/quantifier was 0.338 and the mean %CV was 9.9%.

**Figure 3.**
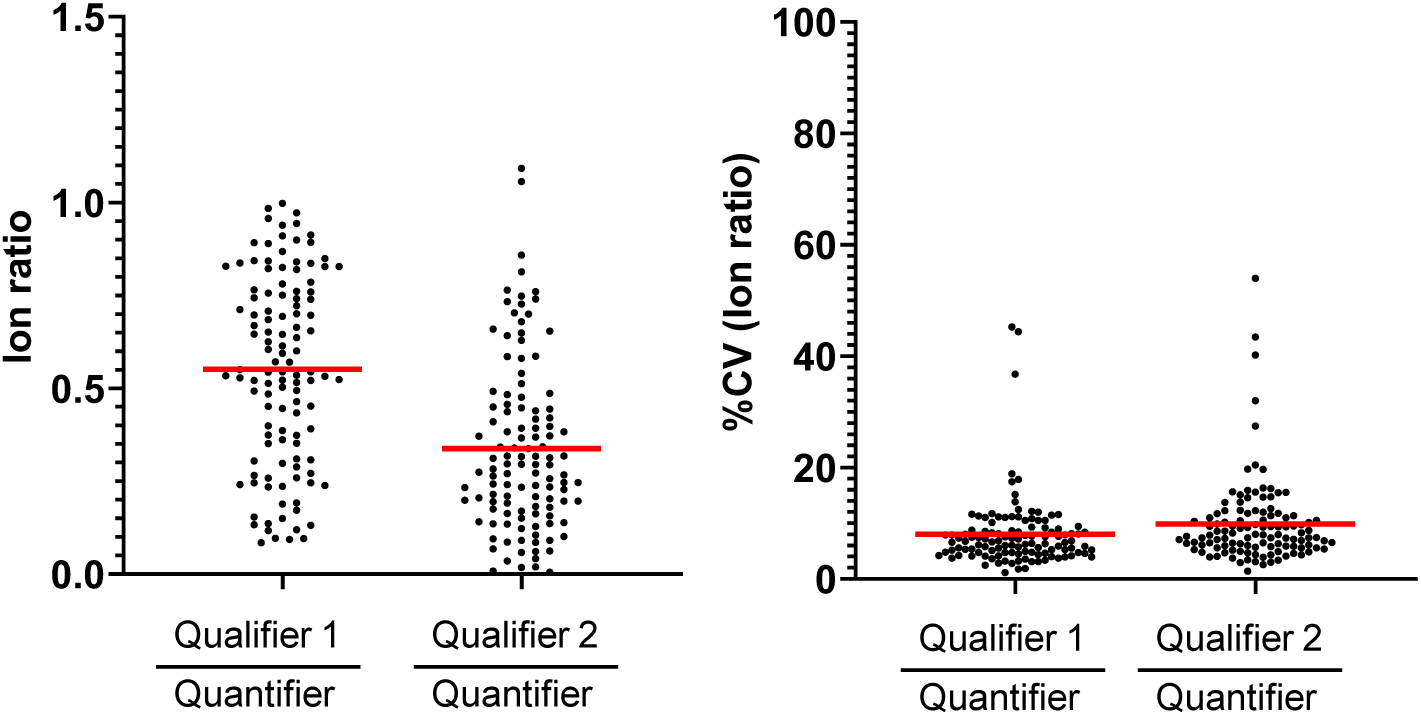
Ion ratios and %CV for 117 HSP peptides in 18 plasma samples.

### Linearity of the HSP assay in plasma matrix

The conventional method to evaluate linearity is to spike serially diluted standard proteins and constant amounts of SIL internal standards into an analyte-free sample matrix and then quantify their peaks to determine the recovery, upper limit of quantitation (ULOQ) and lower limit of quantitation (LLOQ). This was not feasible for the HSP assay because plasma contains endogenous HSP proteins. As an alternative, linearity was determined using serially diluted SIL peptides spiked into a pooled digested healthy plasma matrix (23) (Figure 4). All SIL peptides had a linear response over a concentration range (LLOQ to ULOQ) of at least 100-fold, with linearity confirmed by correlation coefficients (R^2^) of at least 0.94.

**Figure 4.**
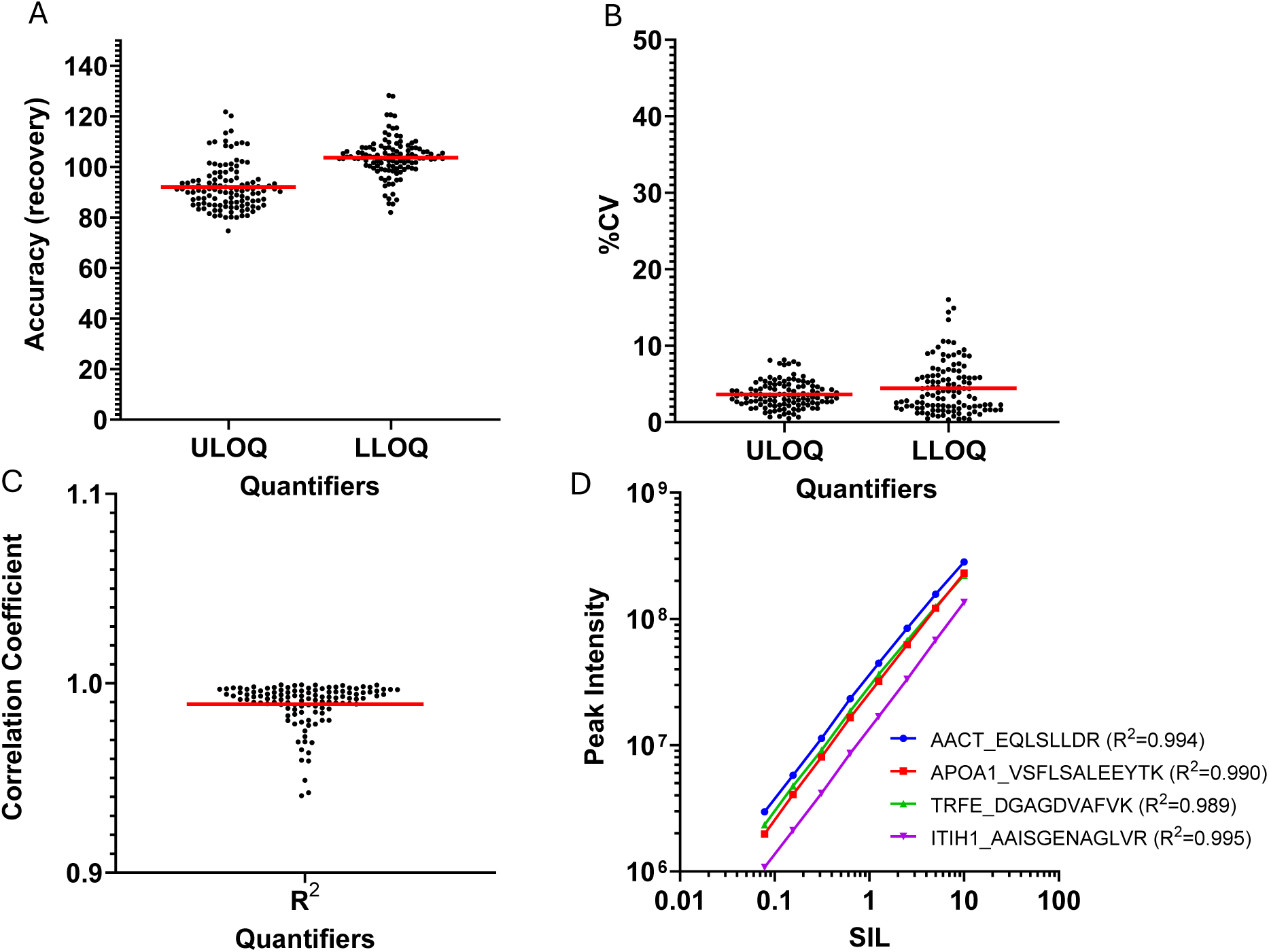
Quantification of SIL peptides diluted into a trypsin digest of pooled healthy plasma.

### HSP assay reproducibility and peptide stability

To measure reproducibility, aliquots of SIL peptides spiked into a trypsin digest of healthy human plasma were injected into the LC-MS/MS 6 times daily for 5 days (one aliquot per day). The peak intensities of the quantifier transitions of the 117 SIL peptides spanned at 6550-fold range (Figure S2A). All quantifier transitions demonstrated excellent signal reproducibility over five days, with average %CVs ranging from 1.5% to 7.0% (Figure S2B). The %CVs trended higher for lower intensity quantifier peptide transitions (Figure S2C).

Peptide stability was evaluated for SIL peptides stored at 14°C in a digested plasma matrix in an LC autosampler. SIL peptide recovery was measured in triplicate every 24 hours for 4 days.

Recoveries for all but one of the 117 SIL peptides varied by <20% over 4 days. Recovery of the 117 peptide varied by <20% for 3 days (Figure S3).

### Identification of CKD biomarkers

The CKD samples came from cases with a rapid GFR decline over three years and were compared to age, sex, race and metabolic disease matched controls (Table 1). Analysis with the HSP assay revealed reduced expression of alpha 2-antiplasmin (*p* = 0.0182 - 0.0353), immunoglobulin heavy constant alpha 1 (*p* = 0.014 - 0.0245), and antithrombin III (*p* = 0.033 - 0.045) in the cases but not in the controls (*p* = 0.5-0.9). ANT3 and IGHA1 were each represented by two peptides and A2AP was represented by three peptides. Reduced expression was observed for all 7 peptides. Data for alpha 2-antiplasmin shows a reduced concentration over time in most CKD cases (Figure 5) and confirms the linearity, reproducibility and accuracy of the measurements (Figure 6). Similar results were obtained for immunoglobulin heavy constant alpha 1 and antithrombin III (Figure S4).

**Figure 5.**
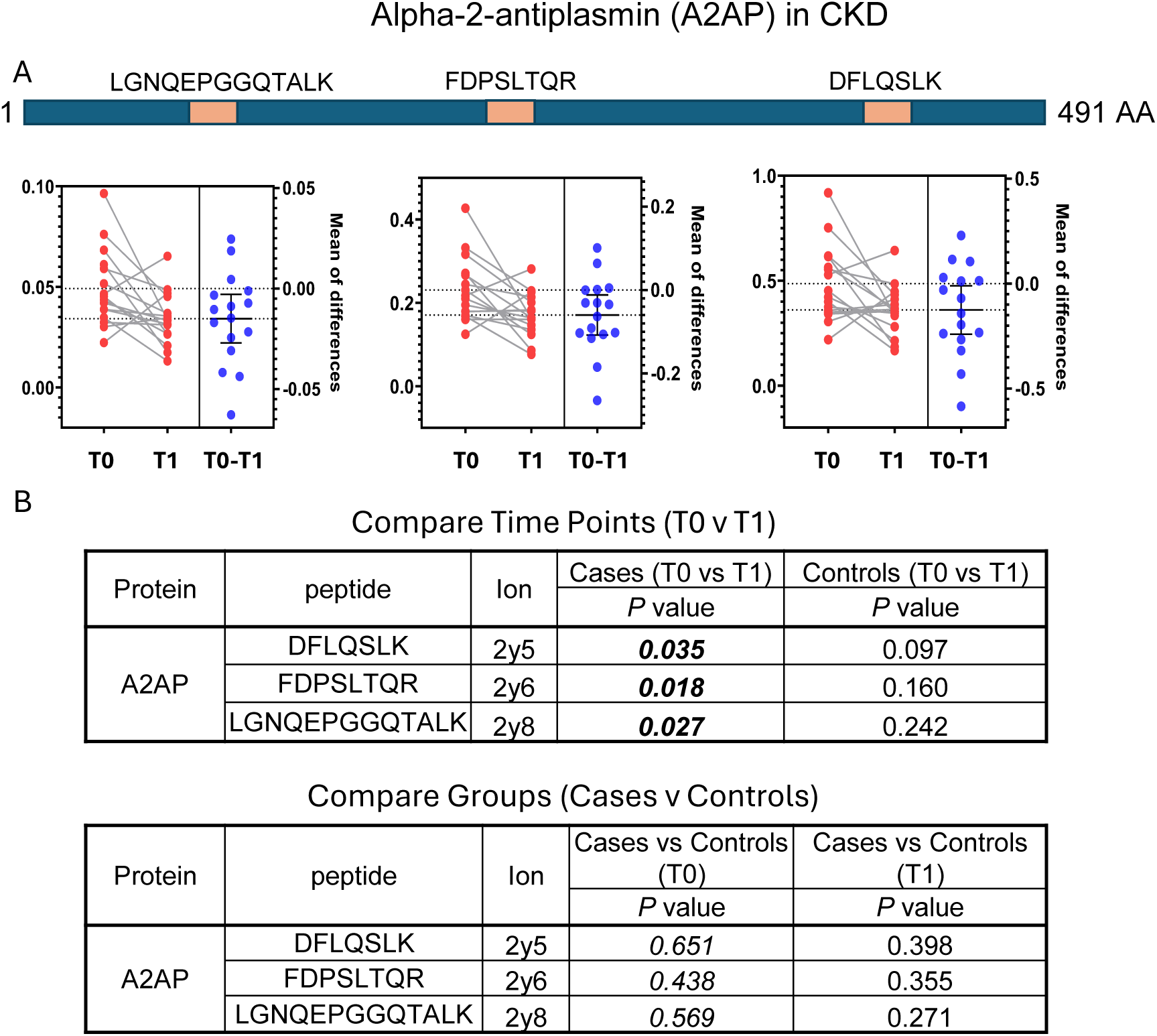
Reduced expression of Alpha-2-antiplasmin (A2AP) correlates with Chronic Kidney Disease (CKD). (A) Three A2AP proteotypic peptides were quantified in samples at the baseline (T0) and final (T1) timepoints. The graphs present means of measurements from diseased individuals (red) and show the change over time (blue). (B) Paired T tests reveal a statistically significant reduction in A2AP at the T1 timepoint in individuals with CKD.

**Figure 6.**
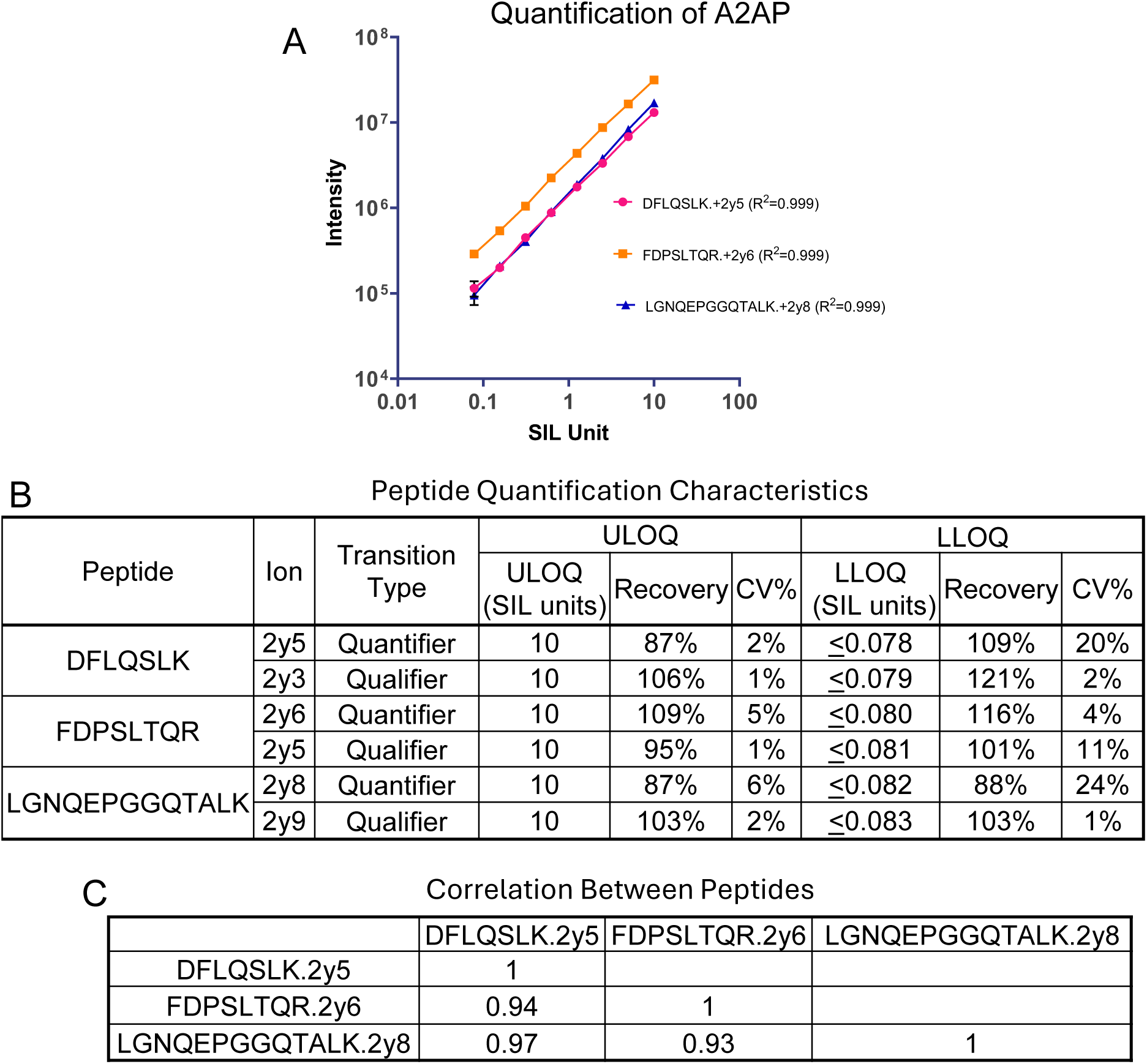
(A) Linearity of SIL peptide quantitation in plasma matrix. (B) Upper and lower limits of quantification of quantifier and qualifier transitions from the SIL peptides. (CF) Correlations between peak areas of the three A2AP peptides in 64 CDK cohort plasma samples.

## DISCUSSION

This report describes an HSP assay that measures the concentration of 60 biologically important plasma proteins which was developed to monitor the health and/or disease state of individuals. We recently reported that the same 60 proteins are also detectible in dried blood spots(24). The HSP assay is a scheduled peptide-MRM method with excellent linearity and reproducibility that targets pre-defined quantifier and qualifier transitions of tryptic peptides from plasma proteins and SIL internal standards. Peptide ion ratios between quantifier and two qualifier ions from the analyte and SIL controls are monitored to ensure data quality.

As a test case, we analyzed plasma samples from the CRIC study, which revealed an association between CKD and reduced expression of alpha 2-antiplasmin, antithrombin-III, and immunoglobulin heavy constant alpha 1. These proteins have been previously linked to kidney disease, but this is first report correlating reduced expression to a reduced GFR.

Alpha 2-antiplasmin (A2AP), a 70 kDa serine protease inhibitor, is a major physiological inhibitor of plasmin (a serine protease) and plays important roles in fibrinolysis(25–27). Kidneys are major sites of A2AP mRNA accumulation in mouse models, specifically in the epithelial cells in the cortex of a kidney (28). Similarly, the human kidney, also contains high levels of A2-AP mRNA, particularly its cortical region. We hypothesize that A2AP could function as a local regulator of plasmin-mediated extracellular proteolysis in the kidney, wherein reduced local A2AP expression would further accelerate the loss of GFR.

Antithrombin-III (ANT3) is a serine protease inhibitor that regulates the blood coagulation cascade and has anti-inflammatory functions (29). Administration of exogenous ANT3 reduces ischemia-reperfusion injury (IRI) of rat liver and kidney (30, 31). Patients undergoing cardiac surgery have low ANT3 activity and a high incidence of acute kidney injury (32, 33).

Immunoglobulin heavy constant alpha 1 (IGHA1) is the constant region of the immunoglobulin heavy chain of IgA, which plays an important role in immune defenses against infection and excludes foreign antigens. IgA’s are a class of immunoglobulins with distinct structure, localization, and receptor interactions (34). IgA is associated with IgA nephropathy, which may link abnormalities of the IgA system and IgA deposits. The HSP assay revealed that circulating IgA expression is reduced by approximately 20% in individuals with CKD. The mechanism for the reduction is unknown but could involve reduced synthesis or increased degradation.

This study introduces alpha 2-antiplasmin, antithrombin-III, and immunoglobulin heavy constant alpha 1 as candidate biomarkers for CKD progression.

## ABBREVIATIONS

MRM: Multiple reaction monitoring
SRM: selected reaction monitoring
FA: Formic acid
ACN: acetonitrile
CKD: chronic kidney disease
CRIC: Chronic Renal Insufficiency Cohort
SIL: Stable Isotope-Labeled
SPE: solid phase extraction
CAD: coronary artery disease
ULOQ: Upper Limit of Quantification
LLOQ: Lower Limit of Quantification
GFR: Glomerular Filtration Rate
HSP: Health Surveillance Panel
MS: Mass spectrometry.

## ACKNOWLEDGEMENTS

This work was supported by 1 U01 DK124019-01 (Design and Validation of Easy-to-Adopt Mass Spectrometry Assays of Importance to Type I Diabetes), 4U54CA260591-02 (Diversity and Determinants of the Immune-Inflammatory Response to SARS-CoV-2)

Funding for the CRIC Study was obtained under a cooperative agreement from National Institute of Diabetes and Digestive and Kidney Diseases (U01DK060990, U01DK060984, U01DK061022, U01DK061021, U01DK061028, U01DK060980, U01DK060963, U01DK060902 and U24DK060990). In addition, this work was supported in part by: the Perelman School of Medicine at the University of Pennsylvania Clinical and Translational Science Award NIH/NCATS UL1TR000003, Johns Hopkins University UL1 TR-000424, University of Maryland GCRC M01 RR-16500, Clinical and Translational Science Collaborative of Cleveland, UL1TR000439 from the National Center for Advancing Translational Sciences (NCATS) component of the National Institutes of Health and NIH roadmap for Medical Research, Michigan Institute for Clinical and Health Research (MICHR) UL1TR000433, University of Illinois at Chicago CTSA UL1RR029879, Tulane COBRE for Clinical and Translational Research in Cardiometabolic Diseases P20 GM109036, Kaiser Permanente NIH/NCRR UCSF-CTSI UL1 RR-024131, Department of Internal Medicine, University of New Mexico School of Medicine Albuquerque, NM R01DK119199.

## CRIC study investigators

Lawrence J. Appel, MD, MPH.; Jing Chen, MD, MMSc, MSc.; Debbie L Cohen, MD.; Harold I. Feldman, MD, MSCE.; Alan S. Go, MD.; James P. Lash, MD.; Robert G. Nelson, MD, PhD, MS.; Mahboob Rahman, MD.; Panduranga S. Rao, MD.; Vallabh O. Shah, PhD, MS.; Mark L. Unruh, MD, MS.

## DATA AVAILABILITY

All data is contained within the manuscript and supplemental data.

## SUPPLEMENTAL DATA

This article contains supplemental data.

## FOOTNOTES

^#^ (Version 5.9, NCI Build 38, available at https://genealacart.genecards.org/.

## Notes

### Competing Interest Statement

The authors have declared no competing interest.

