## SupplementalMaterial for "A targeted LC MS/MS assay of a health surveillance panel and its application to chronic kidney disease"

**Supplement materials and methods**

**DIA analysis**

DIA-MS was performed on an Eksigent 415 LC system operated in microflow mode linked to a 6600 TripleTOF (Sciex), as previously described (17, Robinson et al. 2020). Digested plasma samples were desalted on an HLB plate, resuspended in 2% ACN, 0.1% FA in water, peptides were pre-loaded with a trap column (ChromXP C18CL 10 × 0.3 mm, 5 μm, 120 Å), and then separated on an analytical column (ChromXP C18CL 150 × 0.3 mm 3 μm 120 Å) at a flow rate of 5 μL/min using a linear gradient of 3−35% mobile phase B (0.1 % FA in AC) for 60 min at 30 °C. The column was washed and re-equilibrated with 35−85% B for 2 min, 85% B for 5 min, and 3% B for 7 min. Source parameters were: gas 1 = 15, gas 2 = 20, curtain gas = 25, source temp=100, and voltage = 5500 V. DIA MS1 scans were acquired using a dwell time of 250 ms in the mass range of 400−1250 m/z at 45 000 fwhm. DIA MS2 scans were acquired in high-sensitivity mode at 15,000 fwhm over the precursor range of 400−1250 m/z with the MS2 range of 100−1800 m/z using 100 variable windows with a dwell time of 30 ms.

**DDA analysis**

DDA-MS was performed on an EASY-nLC 1000 connected to an Orbitrap Elite (Thermo) equipped with an EasySpray ion source, as previously described (Parker, Venkatraman, and Van Eyk 2016). Digested plasma samples were desalted on an HLB plate, eluted in mobile phase A, loaded onto a PepMap100 trap column (Thermo, 75 µm x 2 cm, C18 3 µm 100Å), and separated on a PepMap RSLC C18 analytic column (25 cm x 75 μm ID, 2 μm) using a gradient of 5-15% B for 13 minutes. The column was washed and re-equilibrated with 15-30% B for 5 minutes, 30-100% B for 2 minutes, and 100% B for 5 minutes. Full scans were acquired in the Orbitrap with a resolution of 60,000 Hz from 400-1600 m/z. MS2 scans were acquired in the ion trap for the top 15 ions in CID. Charge state screening was enabled to reject unassigned and singly charged ions. Dynamic exclusion was set to a repeat count of 1, repeat duration of 50 seconds, and an exclusion duration of 90 seconds.
