## SupplementalFigures for "A targeted LC MS/MS assay of a health surveillance panel and its application to chronic kidney disease"

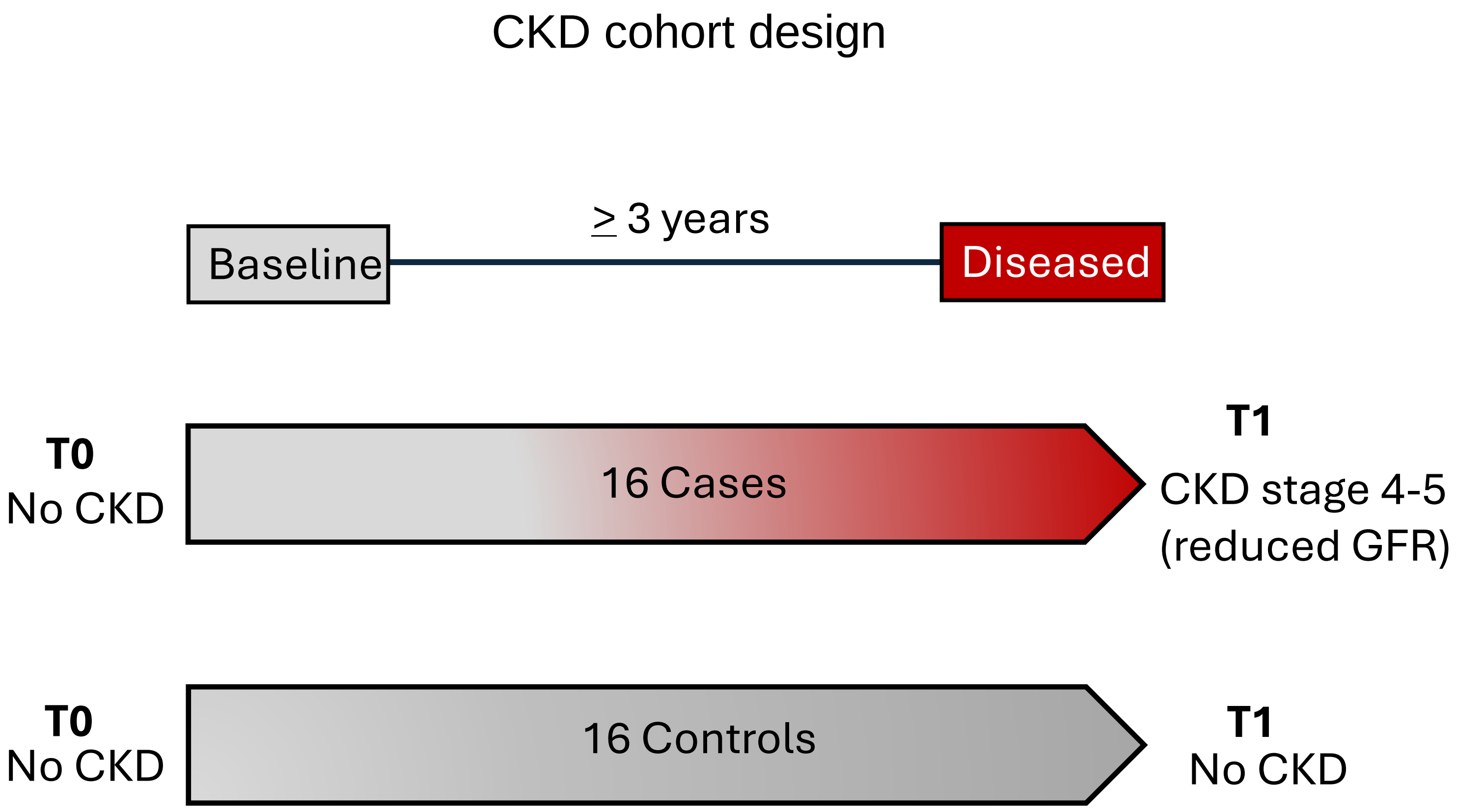


**FigureS1**. CKD cohort design.

### LC-MS/MS reproducibility

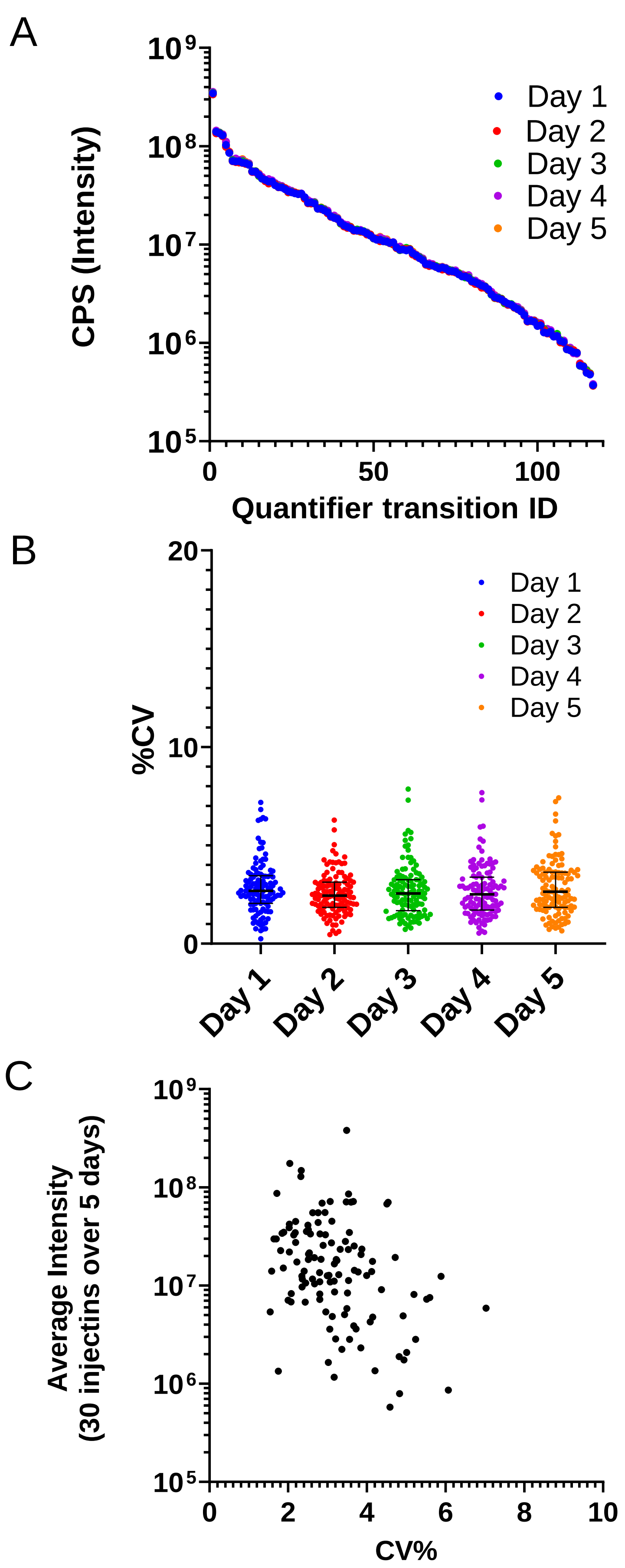


**FigureS2**. LC-MS/MS reproducibility. SIL peptides in digested control pooled plasma were injected 6 times per day on 5 separate days. A) Average intensities. B) Intra-day %CV. C) Inter-day %CV versus average intensities.

SIL peptide stability on the autosampler at 14°C

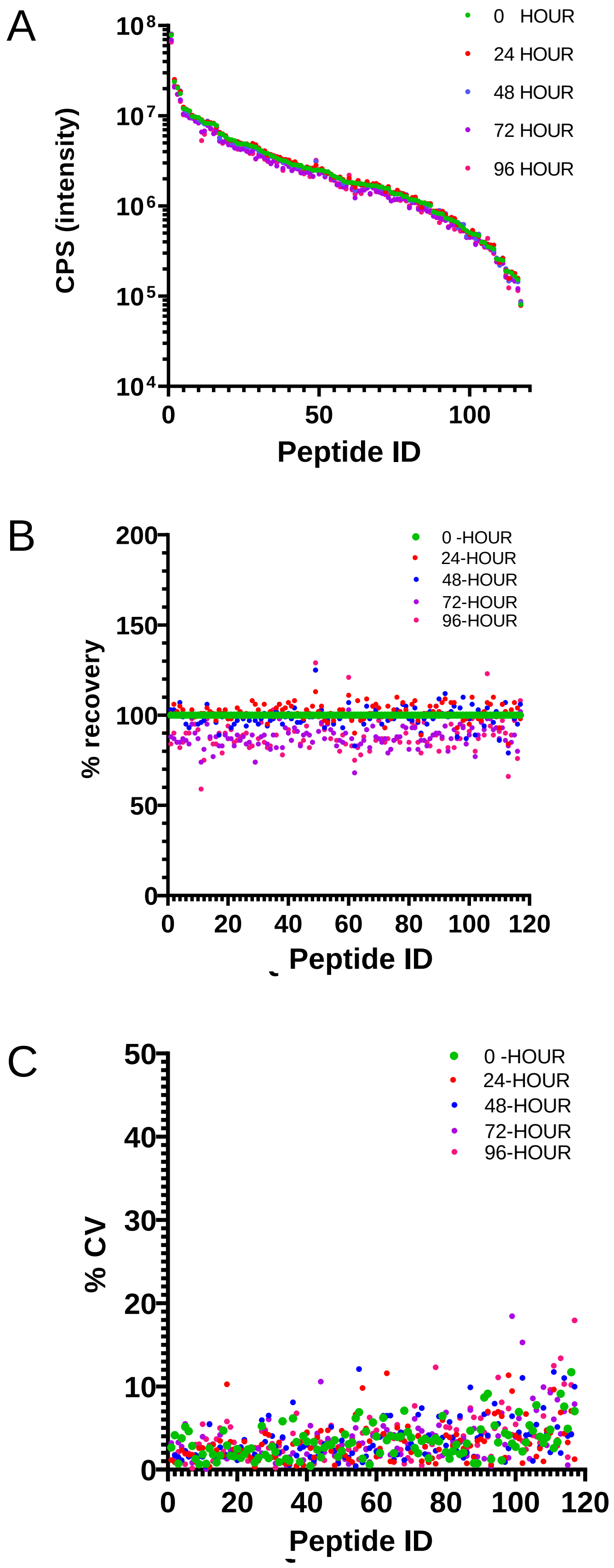


**FigureS3**. SIL peptides were stored at 14 ºC in the autosampler of the HPLC. At 24-hour intervals, three injections were taken from the same vial to quantify 117 peptides and determine intensity (A), recovery (B), and %CV (C).

### Antithrombin-III (ANT3) and Immunoglobulin heavy constant alpha 1 (IGHA1) in CDK

A

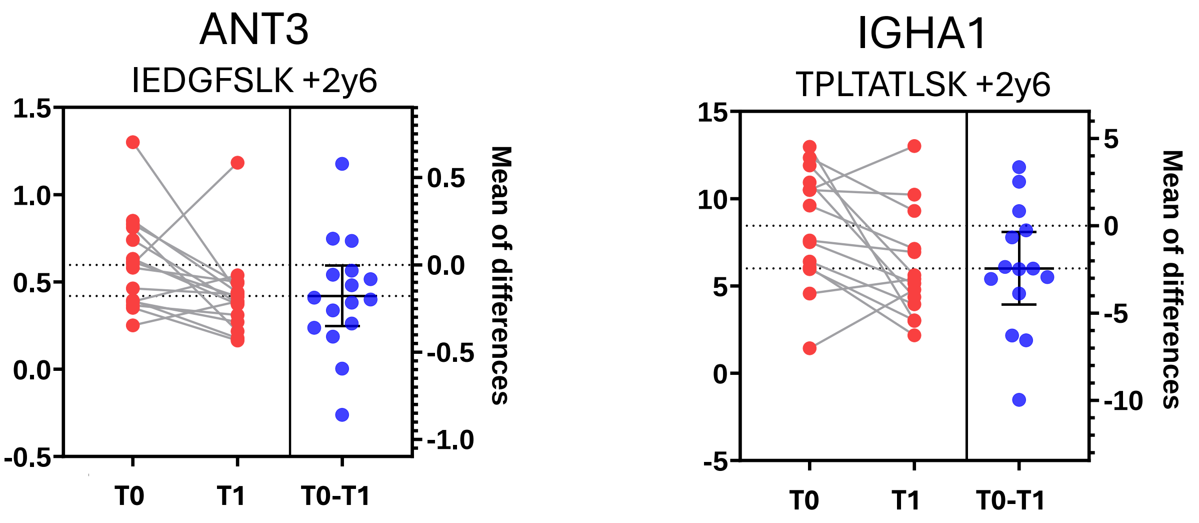


B

Compare Time Points (T0 vs T1)

| Protein | Peptide | Ion | Cases (T0 vs T1) | Controls (T0 vs T1) |
|---|---|---|---|---|
| | | | P value | P value |
| | | 2y6 | ***0.045*** | 0.865 |
| ANT3 | IEDGFSLK | 2y7 | ***0.033*** | 0.901 |
| | | 2y6 | ***0.024*** | 0.569 |
| IGHA1 | TPLTATLSK | 2y7 | ***0.014*** | 0.541 |

Compare Groups (Case vs control)

| Protein | Peptide | Ion | Cases vs Controls (T0) | Cases vs Controls (T1) |
|---|---|---|---|---|
| | | | P value | P value |
| | | 2y6 | *0.062* | 0.840 |
| ANT3 | IEDGFSLK | 2y7 | *0.057* | 0.848 |
| | | 2y6 | *0.866* | 0.223 |
| IGHA1 | TPLTATLSK | 2y7 | *0.919* | 0.233 |

**FigureS4**. Reduced expression of Antithrombin-III (ANT3) and Immunoglobulin heavy constant alpha 1 (IGHA1) correlates with Chronic Kidney Disease (CKD). (A) ANT3 and IGHA1 proteotypic peptides were quantified in samples at the baseline (T0) and final (T1) timepoints. The graphs present means of measurements from diseased individuals (red) and show the change over time with 95% confidence intervals for the mean (blue). (B) Paired T tests reveal a statistically significant reduction in ANT3 and IGHA1 at the T1 timepoint in individuals with CKD.

Quantification of ANT3 and IGHA1

A

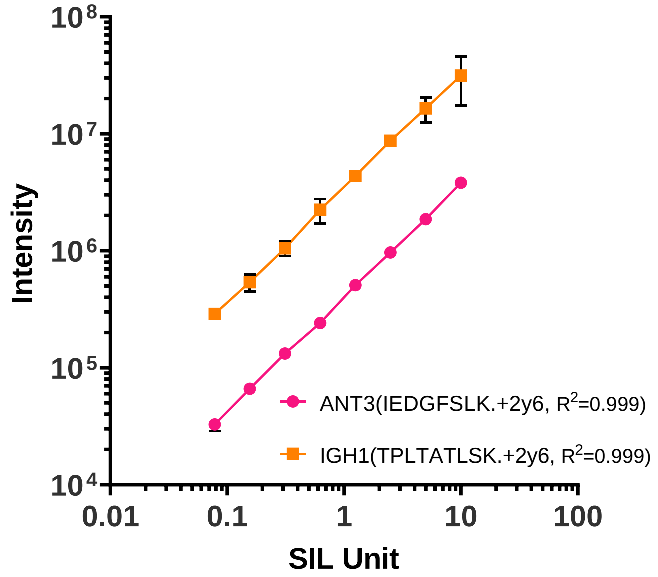


B

Peptide Quantification Characteristics

| Protein | Peptide | Ion | Transition type | ULOQ | | | LLOQ | | |
|---|---|---|---|---|---|---|---|---|---|
| | | | | ULOQ (SIL unit) | recovery | CV% | LLOQ (SIL unit) | Recovery (LLOQ) | CV% (LLOQ) |
| ANT3 | IEDGFSLK | +2y6 | Quantifier | 10 | 100% | 2% | ≤ 0.0780 | 100% | 12% |
| | | +2y7 | Qualifier | 10 | 91% | 4% | ≤ 0.0781 | 91% | 12% |
| IGHA1 | TPLTATLSK | +2y6 | Quantifier | 5 | 87% | 4% | ≤ 0.0782 | 87% | 1% |
| | | +2y7 | Qualifier | 5 | 90% | 5% | ≤ 0.0783 | 90% | 6% |

**Figure 5S**. (A) Linearity of SIL peptide quantitation in plasma matrix. (B) Upper and lower limits of quantification of quantifier and qualifier transitions from the SIL peptides.
